# Investigating post-breeding moult locations and diets of Common Guillemots (*Uria aalge*) in the North Sea using stable isotope analyses

**DOI:** 10.1101/2020.09.01.276857

**Authors:** Alec P. Christie

## Abstract

Seabird movements and diet during the non-breeding season are poorly studied, yet understanding these aspects of seabird ecology is extremely important to effectively conserve these protected species. Stable isotope analyses (SIA) provide a cost-effective solution to filling these knowledge gaps, yielding information on diet and foraging locations of animals. This study aimed to use SIA to investigate whether Common Guillemots (*Uria aalge*) from different age classes and locations in the UK had contrasting diets and foraging areas during the post-breeding moult (July-September). SIA of secondary feathers and a newly-developed North Sea isoscape were used to identify the likeliest foraging areas and diets of deceased guillemots recovered from beaches in eastern Scotland and mixed fisheries in Cornwall and the Celtic Sea. Overall, guillemots foraged widely in the western, eastern and southern North Sea, consuming a variety of clupeid, gadoid and invertebrate prey. There were negligible dietary differences between age classes and birds from different recovery locations. Juveniles showed a wider range in foraging areas, but both age classes foraged in similar parts of the North Sea. Guillemots recovered from Scotland may have foraged further north, only overlapping with guillemots recovered from the southwestern UK in the southern and eastern North Sea. Their winter recovery locations also implied that they exhibited different movement strategies during the non-breeding season, meriting further investigation. Conservation efforts should target foraging areas in the southern and eastern North Sea which are highly threatened by gillnet fishing, shipping traffic and oil infrastructure.

## 1. Introduction

The movements and diets of seabirds during the non-breeding season remain poorly understood aspects of their ecology (Harris et al., 2015). Long-term research has focused on accessible breeding colonies, but once seabirds leave in late summer, they do not usually return until the next breeding season or for occasional visits in autumn (Wanless and Harris, 1986). A better understanding of where seabirds go and what they eat during this time can aid efforts to successfully protect and conserve their populations.

During the non-breeding season, most seabirds undergo one or more post-breeding moults to varying extents. The Common Guillemot (*Uria aalge*) is one example, undergoing a complete flightless moult from July-September as soon as they leave their breeding colonies (Cramp, 1985; Pyle, 2009). During this period of the year they are less mobile as they regrow new feathers at sea (Birkhead and Taylor, 1977) and therefore probably forage within specific areas that may be at risk from human activities – such as bycatch from gillnet fisheries. Various studies suggest guillemots may disperse widely outside of the breeding season (Mead, 1974; Swann and Ramsay, 1983; Reynolds et al., 2011; Harris et al., 2015), possibly with juveniles and immature birds (one to four years old) ranging further from their natal colony than adults (over five years old) (Harris and Swann, 2002). However, most of this knowledge is based on ringing studies from a small number of colonies with limited sample sizes and inherent biases – for example, birds staying close to shore are more likely to be re-sighted, so knowledge of at-sea distributions is poor (Harris et al. 2015).

Within the breeding season, Wright and Begg (1997) suggested distributions of guillemots and their major prey, sandeels (*Ammodytes marinus),* are correlated. However, in the non-breeding season, there is a weaker correlation as their diet includes more sprat (*Sprattus sprattus*) and small herring (*Clupea harengus*). Limited studies on the winter diet of guillemots in the southeastern and western North Sea have shown this to be even more varied, including non-fish prey items such as molluscs, crustaceans (crabs) and polychaete worms, in addition to a high proportion of clupeids, sandeels, pipefishes and gobies (Lorentsen and Anker-Nilssen, 1999; Sonntag and Hüppop, 2005). Blake et al. (1985) found that a seasonal switch-over occurs in guillemot diet in the northern North Sea, as from March-August they mainly consume sandeels, but from September into winter they forage on clupeid and gadoids. The post-breeding moult of guillemots bridges these two periods, making it a time of major dietary changes. However, no studies have yet considered whether adults and juveniles consume different prey during this period, or whether these dietary changes in the post-breeding moult also occur further south in the North Sea. Understanding guillemot diet during this time is particularly important to evaluate whether they come into conflict with fisheries, which arguably pose the greatest threat to guillemot populations (Wright and Begg, 1997; Peterz and Blomqvist, 2010). Moreover, knowing where guillemots travel to carry out the post-breeding moult can help ascertain whether other human activities, such as oil spills (Le Rest et al., 2016), pose a significant risk to their moulting aggregations.

With 13% of the global Common Guillemot population and one-third of the North Atlantic population concentrating in European shelf seas during the non-breeding season (Mitchell et al., 2004; JNCC, 2016; Le Rest et al., 2016), filling these knowledge gaps becomes even more important to the conservation of this seabird. In the northern hemisphere, these guillemots are also considered one of the most important fish-eating seabirds by biomass. Therefore, further investigation into how anthropogenic exploitation and natural variation affect their populations is vital to better understand the North Sea ecosystem and how its fish stocks are regulated (Lorentsen and Anker-Nilssen, 1999). Furthermore, Common Guillemots are amber-listed as Birds of Conservation Concern (Eaton et al., 2015), making their protection mandatory under the Marine Strategy Framework Directive (EU, 2008).

The main aim of this study was to determine where Common Guillemots carry out their post-breeding moult. More specifically, this study aimed to investigate if adult and juvenile guillemots forage in different locations during this time. Hypothetically, adult and juvenile foraging areas should be relatively similar, since a parental guardian (usually the male) accompanies their offspring out to sea at the end of the breeding season – although the length of time adults stay with juveniles is unknown (Jones and Rees, 1985; Wanless and Harris, 1986). However, juveniles do not return to their natal colony until their second year and are also thought to disperse more widely than adults in the non-breeding season (Harris and Swann, 2002). Moreover, non-guardian birds (usually breeding females and non-breeders) may forage in other locations; females may stay closer to their breeding colony so they can return in October to secure nest sites for the next breeding season (Wanless and Harris, 1986; Halley et al., 1995). Considering sex-related differences in foraging locations would be useful, but unfortunately limited resources meant this was not possible in this study.

This study also aimed to investigate whether birds recovered from two different locations: (1.) ‘southern’ birds from mixed static net Cornish and Celtic Sea fisheries, in southwestern England; and 2.) ‘northern’ birds from beaches in St Andrews, in eastern Scotland foraged in different areas during the post-breeding moult. Distance to colony has been found to be a major factor during the breeding season as guillemots are central place foragers (Wright and Begg, 1997). However, in the non-breeding season they are less constrained by colony location – thus, recovery location may not imply colony location. Nevertheless, differences might be expected between birds recovered from different locations as they may use different movement strategies during the post-breeding moult and the wider non-breeding season. These movements can be tentatively inferred from their different winter recovery locations and post-breeding moult foraging areas. McFarlane Tranquilla et al. (2014) found differences between guillemots in their winter movement strategies in the northwest Atlantic, so it is possible guillemots from the North Sea also exhibit different movement strategies.

This study also aimed to investigate the diet of guillemots during the post-breeding moult, as well as any dietary differences between age classes and recovery locations. Based on previous studies of guillemot diet in the non-breeding season (Blake et al., 1985; Lorentsen and Anker-Nilssen, 1999; Sonntag and Hüppop, 2005), adult diet during the post-breeding moult should be composed of a mixture of different fish and non-fish prey discussed earlier. Few studies have considered if there might be differences between adult and juvenile diet during the non-breeding season. Lorentsen and Anker-Nilssen (1999) suggested that adults and juveniles may eat the same range of prey but have different preferences during winter, with adults preferring gadoids and juveniles preferring clupeids, gobids and polychaetes. These preferences were explained by age-related differences in swimming ability, hunting experience and body condition (Lorentsen and Anker-Nilssen, 1999). Hypothetically, these patterns should also occur during the post-breeding moult. Moreover, diet should not vary between age classes or recovery locations if birds forage in similar locations during this period. However, this could differ if they foraged at different latitudes where switch-overs in diet discussed earlier might occur at different times in the year due to seasonal cues or prey availability.

To date, seabird movements and diet outside of the breeding season have been investigated using several methods. Traditionally, stomach contents analyses were undertaken to investigate diet and infer foraging locations from the distribution of identified prey (Votier et al., 2001). More recently, with decreasing size and weight of GPS satellite tags and light-based geolocators (using day length and sunrise/sunset timings to determine longitude and latitude), tagging of seabirds has been undertaken to track movements (Bridge et al., 2013). However, GPS tags are still too large and heavy for many seabird species, whilst lighter, smaller geolocators suffer from inaccuracy at high latitudes (Harris et al., 2015) and during equinoxes (Hebblewhite and Haydon, 2010; Bridge et al., 2013). Marine isotope-based geographic assignment is a relatively recent development in stable isotope analyses (SIA) which offers the potential to investigate both diet and foraging location using a single method (MacKenzie et al., 2014; Trueman et al., 2016).

As a biogeochemical approach, SIA exploits the fact that the isotopic composition of prey items (and ultimately the primary productivity in the environment) is incorporated into consumer tissues during tissue synthesis and growth. Therefore, tissues that become metabolically inert after their growth period (such as feathers) contain isotopic signatures that can help isolate the diet and foraging areas of an animal during specific periods of the year (Hobson and Bond, 2012). Trueman et al. (2016) recently built a marine isoscape (‘isotopic map’) for the North Sea using spatial variability in carbon and nitrogen isotopes measured from jellyfish tissues. This reflects spatial variation in primary production (phytoplankton) and allows isotopic signatures from animal tissues to be matched to areas with similar isotopic compositions to infer foraging locations. Trueman et al. (2016) successfully assigned scallops and herring to regions within the North Sea with accuracy and precision that rivals GLS tagging methods – 200-300km for SIA compared to 200-400km for GLS (at lower latitudes and outside of equinoxes; Phillips et al., 2004; Lisovski et al., 2012). However, whilst several studies using tissues from known locations have supported the validity and robustness of isotope-based assignment (Caccamise et al., 2000; Wassenaar and Hobson, 2000; Wunder et al., 2005), this method has a trade-off between accuracy and precision. This is illustrated by Trueman et al. (2016), in which assignments to regions of 50% of the North Sea were 100% accurate, but assignments to regions 30% of the North Sea (a scale of ∼10^5^ km^2^) were only 75% accurate. Despite this, however, isotope-based geographic assignment still offers a valid alternative to GLS tagging methods. Furthermore, SIA has several advantages over GLS in relation to researching seabird movements and diet during the non-breeding season. One such benefit is greater cost-effectiveness through attaining larger sample sizes with limited funding. GLS would otherwise require expensive, invasive tagging and recapturing – the latter of which may not be possible (Hebblewhite and Haydon, 2010; Bridge et al., 2013). Due to constraints in the budget for this study, SIA was therefore the optimal method to use.

Nevertheless, it is important to acknowledge several caveats before valid conclusions can be drawn from isotope-based geographic assignment. Firstly, isotopic values reflect the origin of primary productivity that fuel higher trophic levels (Trueman et al., 2016), not the individual location of an organism. Therefore, when considering data for animals from upper trophic levels, with high mobility, and relatively slow isotopic assimilation rates, the ecological meaning may be harder to interpret (Trueman et al., 2016). This is because it is possible that animals or their prey move away from the origin of primary productivity before assimilation of isotopes to higher trophic levels occurs. Therefore, there is a balance between animal mobility and isotopic assimilation rate that determines if isotope values reflect the true foraging location of an upper trophic level consumer during a certain period. However, these assumptive problems can be resolved by choosing the correct tissues for analysis. Feathers can allow more accurate geographic assignment of highly mobile birds to foraging areas since they are grown and assimilate stable isotope ratios during specific times of year: moults (Jaspers et al., 2004; Quinn et al., 2016). This is because assimilation in feathers ceases after a feather is fully grown due to atrophy (Burger and Gochfeld, 2000), thus isolating the origin of primary productivity during the moult period when feathers grow. Additionally, since the moult is a flightless period, birds are less mobile and therefore their foraging location is more likely to be the same as the origin of assimilated primary productivity. Using knowledge of guillemot moulting chronology, secondary feathers were selected as these are grown during the post-breeding moult (Harris and Wanless, 1990) and therefore their isotopic compositions should reflect where guillemots foraged and what they consumed during this time. However, it should also be acknowledged that guillemots recovered from fisheries are likely to be less biased samples, since they were probably healthy before being killed as bycatch (Peterz and Blomqvist, 2010). Conversely, guillemots recovered from beaches may have been sick or injured birds and moved abnormally before they stranded, making them less representative samples. Therefore, age-related differences were also considered within these different recovery locations to check and account for these different levels of bias.

Using the isoscape developed by Trueman et al. (2016), this study aimed to use stable isotope analyses of feathers to compare the North Sea foraging areas and diet of Common Guillemots from different age classes and recovery locations during the post-breeding moult. According to extensive searches of the literature, no studies have yet attempted to investigate these differences with this approach.

## 2. Materials and methods

### 2.1. Field collection

14 carcasses of Common Guillemots (*Uria aalge*) were collected from East Sands, Aquarium Sands and West Sands in St Andrews, Scotland between September-October 2016 (Appendix 6.1-6.3). Another seven carcasses were retrieved by fisheries observers from mixed static net fisheries in Cornish waters and adjacent Celtic Sea waters between December 2016 and February 2017, making 21 birds in total. Guillemots recovered from these southwestern fisheries are termed ‘southern’ birds, whilst those recovered from Scotland are termed ‘northern’ birds, referring to their last known geographical location.

### 2.2. Age determination

21 Common Guillemots were separated into two age classes: 1.) first-years and 2.) second-years and older by inspecting greater underwing covert feathers. White tips distinguish first-year birds from older birds which possess dark tips (Kuschert et al., 1981). Distinguishing immature birds (two to four years old) by presence of white tips and a cloacal bursa was impossible as most carcasses only possessed wings (Kuschert et al., 1981). Within this study first-year birds were termed ‘juveniles’, whilst second-year and older birds were termed ‘adults’ – but may have included immature and mature birds.

### 2.3. Stable isotope preparation

Birds were frozen as soon as possible after collection and stored in sealed plastic bags. Carcasses were thawed to cut off two randomly-selected secondary feathers at two-thirds of the way to shaft’s base. To remove dirt, feathers were cleaned with warm tap water and then an ultrasonic bath for 10 minutes. Decontamination of feathers was performed using three rinses with deionised water and a 2:1 DCM:Methanol solution. A final rinse with deionised water was undertaken and feathers were fully dried in an oven at 115°C overnight. Two subsamples (approximately two mm wide) were cut from each feather using decontaminated stainless steel scissors, avoiding the rachis and cutting perpendicular to the barbs. Scissors were cleaned with ethanol and rinsed with deionised water before each subsample was taken to avoid contamination. These replicates were taken to account for isotopic variation within individuals. Subsamples were weighed to 0.5-0.8mg (recorded to three significant figures) and placed into standard tin capsules for stable isotope analyses.

### 2.4. Stable isotope analyses

Stable isotope analyses were undertaken at the Natural Environment Research Council Life Science Mass Spectrometry Facility, East Kilbride (Scotland) using a continuous-flow isotope mass spectrometer and a Flash HT elemental analyser (2012).

Isotopic ratios were given in δ notation in parts per thousand (‰) relative to Pee Dee Belemnite (δ^13^C) or air (δ^15^N) standards. To correct for drift and linearity effects and monitor accuracy and precision, several laboratory standards (including GEL, ALAGEL, GLYGEL and the independent international standard USGS40) were appropriately placed throughout the sample runs. Overall, precision (σ of standards) was ≤0.11‰ for δ^13^C and ≤0.36‰ for δ^15^N values. Accuracy in this laboratory for both δ^13^C and δ^15^N values relative to long-term average values was within 0.12‰ for all four standards.

### 2.5. Isotope-based geographic assignment

#### 2.5.i. Statistical methods for isoscape calibration and assignment

The methodology that follows has been adapted from Trueman et al. (2016) and was carried out by Dr Clive Trueman at the National Oceanography Centre, University of Southampton. This section details the statistical methodology used to build the North Sea isoscape (Fig.1) and how guillemot feathers were calibrated and geographically assigned to this model; the next section (2.5.ii) details how these assignments were displayed graphically.

**Figure 1.**
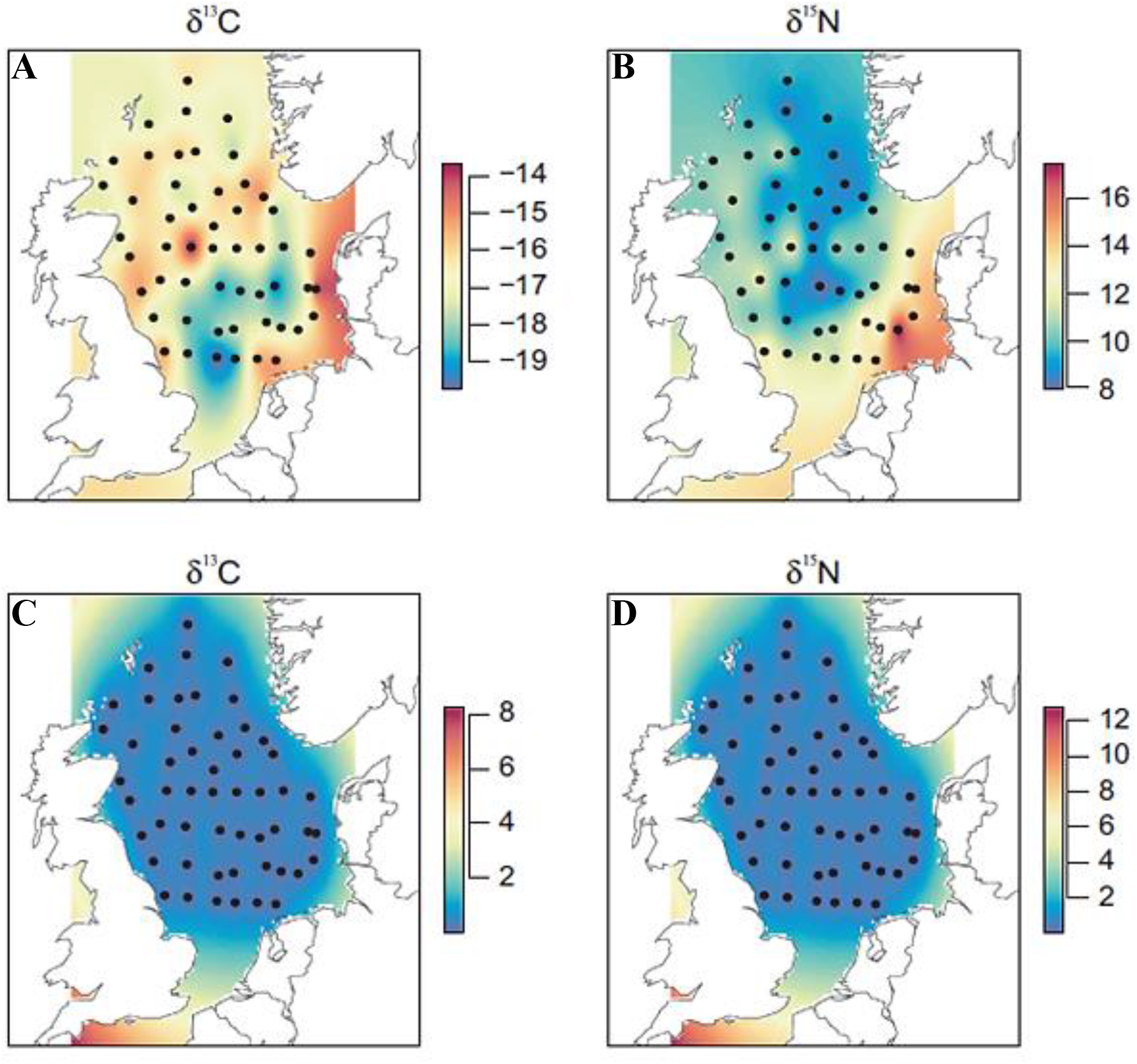
Maps showing the boundaries of North Sea isoscape models (A, B) with associated variances (C, D) for δ^13^C (A, C) and δ^15^N (B, D) values based on jellyfish (Cyanea capillata) sampled in 2011/2015 (adapted from Trueman et al., 2016). The sampling stations for jellyfish are indicated with filled circles. Colours represent δ^13^C and δ^15^N values (A, B) and variances associated with these (C, D) based on the appropriate scales beside each map – red indicates higher values, blue indicates lower values. Extrapolations to areas outside the main sampled area of the North Sea results in greater variance in areas such as the English Channel and limited parts of the UK’s western shore.

Trueman et al. (2016) outline their field methodology for collecting data for their jellyfish-based isoscape model for the North Sea (and limited parts of the UK’s western shores and English Channel). Linear kriging (a common geostatistical interpolation method) was used to generate isoscape models of spatial variation in δ^13^C and δ^15^N values from lipid- and size-corrected isotopic data from jellyfish (*Cyanea capillata*) tissues sampled in 2011/2015.

Isotope-based geographic assignment is based on the probability of a sample originating from a certain location or cell in the isoscape. This in turn depends on the isotopic differences between isotopic values of samples and cells, relative to the total variance in the isoscape. This total variance within the isoscape is composed of two terms: 1.) a spatially varying term drawn from the physical distance between jellyfish sample points estimated using the linear kriging process and 2.) a fixed term reflecting measurement error and between-individual variance (Bowen et al., 2014). Measurement error for jellyfish tissue analyses was determined as the standard deviation from 13 replicate analyses of glutamic acid standards: 0.1‰ for δ^13^C and 0.2‰ for δ^15^N values. Between-individual variance for jellyfish were found (using data from 2011 (MacKenzie et al., 2014) and 2015) to be 1.69‰ for δ^13^C and 1.04‰ for δ^15^N values. Total uncertainty in the isoscape was defined as:

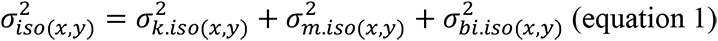

where 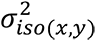 is pooled variance associated with isoscape predictions; 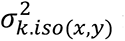 is variance associated with the spatial interpolation model; 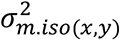 is variance associated with measurement error; and 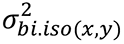 is variance associated with *in situ* between-individual variation.

Measurement error (precision) for guillemot isotope analyses were given in section 2.4., whilst between-individual variances for each isotope and map combination are given below (Table 1).

**Table 1.** Between-individual variance in lipid-corrected δ^13^C and δ^15^N values for all guillemot datasets used to create likely foraging area maps.

| | $\delta^{13}\text{C}$ (‰) | | $\delta^{15}\text{N}$ (‰) | |
| --- | --- | --- | --- | --- |
|  | Juvenile | Adult | Juvenile | Adult |
| **Northern** | 0.46 | 0.002 | 0.67 | 0.0009 |
| **Southern** | 1.69 | 1.90 | 1.59 | 0.27 |
| **Overall** | 0.77 | 0.53 | 0.96 | 0.72 |

These two measures of uncertainty were used to calculate pooled assignment error as follows:

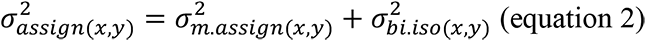

where 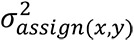 is pooled variance associated with the isoscape prediction; 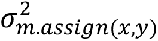 is variance associated with measurement error; and 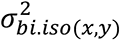 is variance associated with *in situ* between-individual variation.

Calibration between the isoscape model and feathers also comes with uncertainty which was estimated by combining the uncertainty associated with trophic distance and trophic fractionation between jellyfish and guillemots. Although jellyfish sampled in 2015 were of different sizes and weights, this is not expected to cause any size-related differences in their trophic level (Fleming et al., 2015). Therefore, using mass balance (Ecopath) modelling of the North Sea community by Mackinson and Daskalov (2007), a trophic level of 3.6 was assigned to these jellyfish (classed as gelatinous zooplankton). Guillemots were assigned a trophic level of 3.5 (seabirds in Mackinson and Daskalov (2007)), giving a trophic distance of -0.1. A standard deviation of 0.05 was used to ensure 95% of estimates for trophic distance fell between -0.2 and 0 trophic levels (Appendix 6.4). Uncertainty associated with isotopic fractionation between feathers and guillemot prey was estimated as 1‰ for δ^13^C and 3.4‰ for δ^15^N values (a standard deviation of 0.5‰ was given so 95% of estimates fell between 0–2‰ for δ^13^C and 2.4–4.4‰ for δ^15^N values; Appendix 6.4). Bootstrapping 10 000 trophic fractionation and trophic distance values from these distributions then gave estimates of the distribution of isotopic separation values between jellyfish and guillemots (carried through to equation 3).

More uncertainty could be associated with differences in biochemical compositions of jellyfish bell tissues compared to guillemot feathers. Comparing trophic level-corrected guillemot data to the whole range of isotopic values in the isoscape allowed the determination of the smallest offset term required to ensure all guillemot data fell within the range of the isoscape. This minimum tissue-specific adjustment was +2‰ and +0.5‰ for δ^13^C and δ^15^N values, respectively, with standard deviations of ±0.5‰ for both (Appendix 6.4).

Furthermore, this adjustment allowed for possible systematic under/overestimation of trophic differences or isotopic fractionation.

Combining the variance for tissue-specific adjustments with distributions for trophic fractionation and trophic distance separation values gave the estimated variance associated with calibration between guillemot feathers and jellyfish tissues:

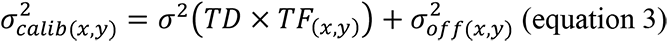

where x and y refer to δ^13^C and δ^15^N values, respectively; TD refers to the distribution of trophic distance values; TF is the distribution of isotopic fractionation values; and 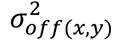 denotes the estimated variance of the tissue-specific adjustment.

The aforementioned variance terms allowed robust assignments of individual birds to the isoscape following an approach set out in Vander Zanden et al. (2015):

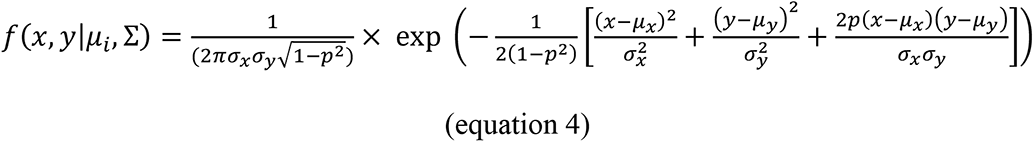

where *f*(*x, y*|*μ_i_*, Σ) is the probability that an individual bird with the according adjusted isotopic composition (where x = δ^13^C and y = δ^15^N) foraged within a given cell (denoted as *i*) in the isoscape; with mean isotopic composition equal to the components of vector *μ_i_* and variance-covariance matrix Σ. *p* is the correlation between δ^13^C and δ^15^N values within the isoscape, whilst *σ_y_* and *σ_x_* are pooled standard deviations for δ^13^C and δ^15^N values, respectively, given below using variances from equations 1-3:

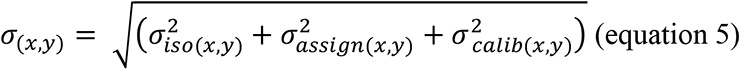

#### 2.5.ii. Presentation of assignment outcomes

Maps displaying assignment outcomes of likely foraging areas for guillemots were produced by Dr Clive Trueman using the following methodology adapted from Trueman et al. (2016). Once statistical assignments were completed, assignment outcomes for guillemots were transferred to a map of the North Sea isoscape. Odds ratios set threshold values that determined the most likely cells in which an individual guillemot foraged during the post-breeding moult (Van Wilgenburg et al., 2012; Vander Zanden et al., 2015). Calculating the odds of a guillemot foraging in a given cell was based on the probability of this occurring relative to the probability of it not occurring (in mathematical notation: *P/1-P);* likelier events have higher odds. An odds ratio can be defined as a ratio between the odds of the outcome taking place, relative to the odds of the most likely outcome based on supplied data:

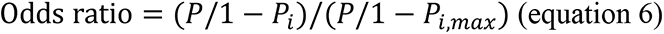

If probability values of cells are greater than this threshold value, then that cell will be defined as a cell of likely foraging area. Taking the reciprocal of the odds ratio gives the total expected proportion of data that falls within the threshold limit, given the normal probability density function. For guillemots, an odds ratio threshold of 1.42:1 was used. This means all cells containing the most likely (1-1.42^-1^) ∼30% of all data outcomes were defined as likely foraging areas, equalling ∼30% of the total isoscape area (Appendix 6.4). The odds ratio threshold therefore determines precision; in this case ∼30% of the North Sea. Trueman et al. (2016) suggests accuracy is ∼75% when using this isoscape to make assignments to foraging areas representing ∼30% of the total area of the North Sea.

Following Van Wilgenburg and Hobson (2011), for every bird, cells determined as likely foraging areas were assigned values of one, whilst all other cells were assigned values of zero. Individual guillemots were pooled and split between age classes (Fig.2A, B) and then split further by location (Fig.2C–F). Summation of values for each cell across the total number of individual birds and division by the total number of birds provided an index for the most frequently assigned cells (presented alongside assignment maps; Fig.2). These maps indicate where guillemots are likely to have foraged during the post-breeding moult, with the numbers of individuals assigned to different cells showing areas of overlap between individuals (darker blue colours mean more individuals were assigned to that area; Fig.2).

**Figure 2.**
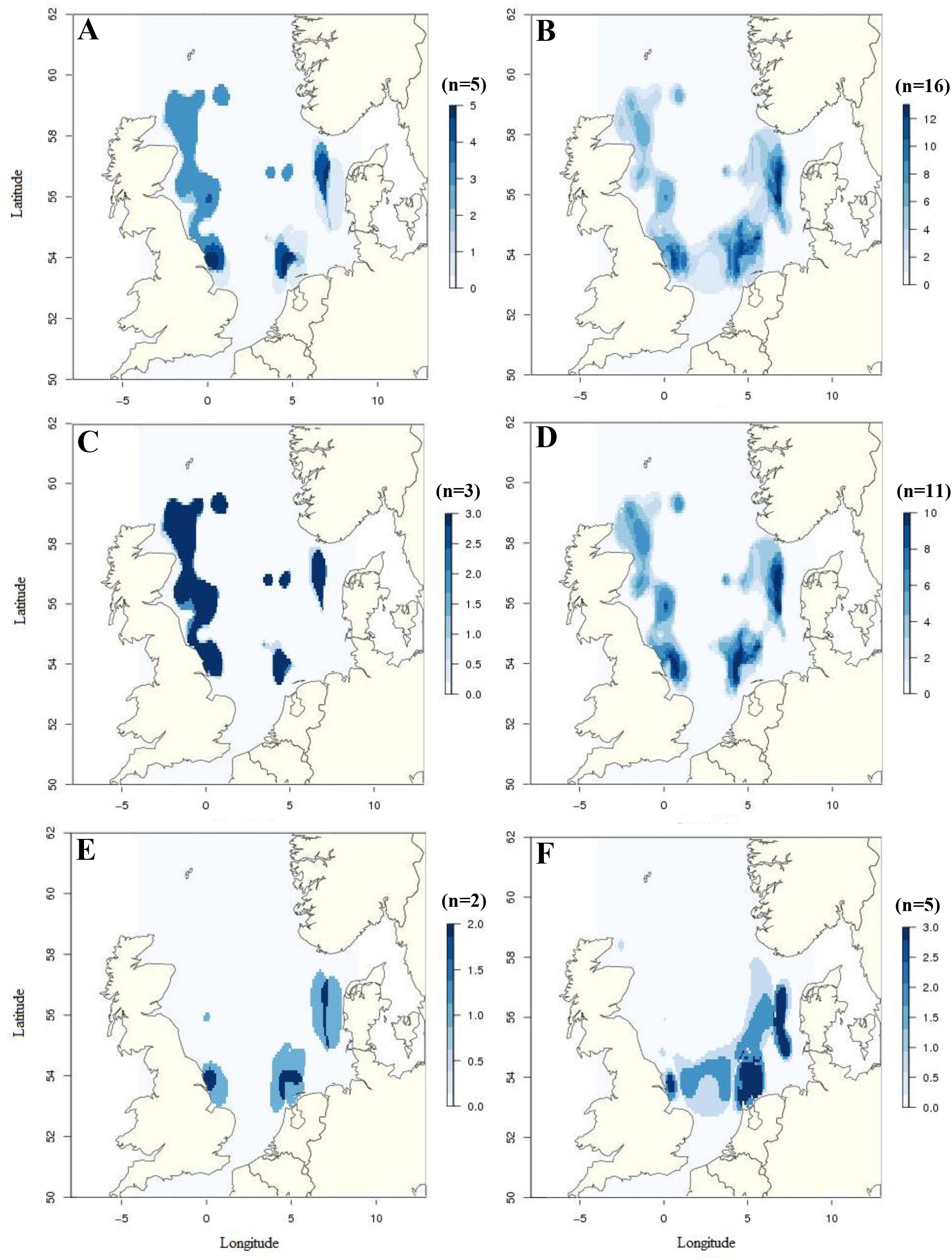
Maps showing most likely foraging locations of common guillemots during the post-breeding moult through isotope-based geographic assignment to isoscape in Trueman et al. (2016). (A) and (B) show foraging locations of adult versus juvenile guillemots, respectively, overall. (C) and (D) show adults versus juveniles for northern birds from St Andrews. (E) and (F) show adults versus juveniles for southern birds from fisheries off southwestern England. Darker blue indicates more birds were assigned to that area, as shown by corresponding legends.

### 2.6. Data analyses

#### 2.6.i. General linear models and cluster analysis

The statistical programme R (R Core Team, 2016) was used to carry out all data analyses and create plots of data. General linear models tested for significant differences in mean δ^13^C and δ^15^N values between age classes, as well as between northern and southern birds. To account for different levels of bias in northern and southern samples, age-related differences were also investigated within these subsets. Model selection was performed using Akaike Information Criteria corrected for small sample sizes (AICc). Normality of residuals, collinearity, leverage effects and homoscedasticity assumptions were satisfied on reviewing residual plots, Shapiro-Wilks Normality and Bartlett’s homogeneity of variance tests.

Cluster analysis was performed using the K-means non-hierarchical clustering function (kmeans) from package stats in R (Afifi and Clark, 1990). This identified the most cohesive clusters possible within the whole guillemot bivariate dataset. To account for weaknesses in the K-means cluster analysis method, 100 initial configurations of selected centroids were run to find the best possible clusters. A plot of total within-groups sums of squares against the number of clusters helped determine the appropriate number of clusters for the method to identify within these data (Appendix 6.5) – levelling off in the line is seen after four clusters, so k=4 clusters were assigned (Fig.3B).

**Figure 3.**
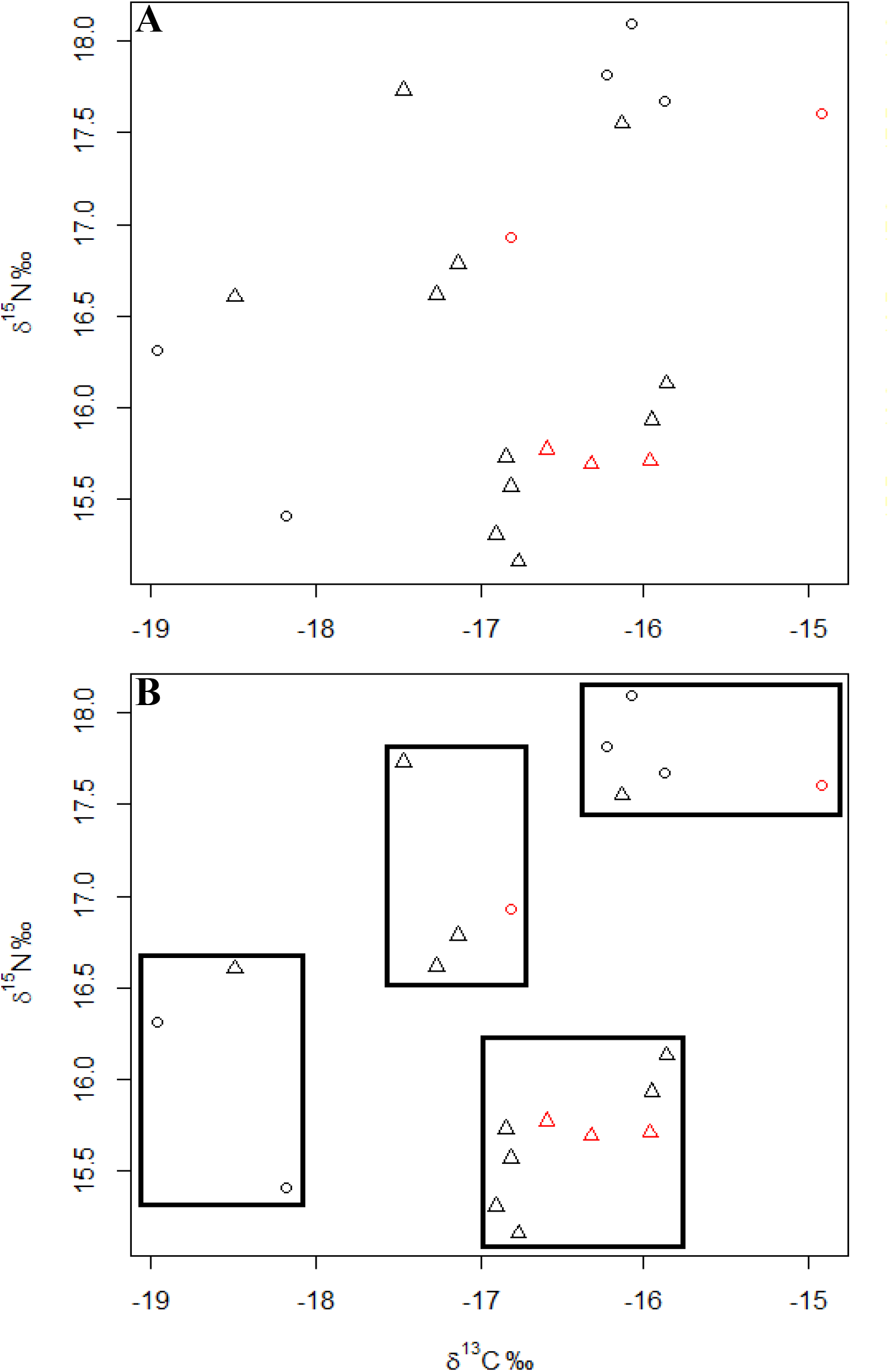
Isotopic biplots showing δ^13^C and δ^15^N values for all birds (A) and when grouped by K-means cluster analysis (B). Triangles denote northern birds, whilst circles denote southern birds. Red symbols are adults, whilst black symbols are juveniles. Boxes contain the four clusters determined by K-means cluster analysis.

#### 2.6.ii. Dietary analyses

To examine diet during the post-breeding moult, trophic enrichment factors (TEFs) were used to compare isotopic values between guillemot feathers and possible prey from the North Sea (using data for fishes from Käkelä et al. (2007) and invertebrates from Kürten et al. (2013); Appendix 6.6). The same TEFs between feathers and prey muscle and associated standard deviations were used as in Meier et al. (2017): 1.9‰ (±0.5‰) for δ^13^C and 3.7 (±1‰) for δ^15^N values. This allowed comparisons between the ranges of predicted prey isotope values for different groups of guillemots and measured mean prey isotope values with associated standard deviations (Fig.4). Temporal stability in the isotopic composition of the North Sea over at least decadal timescales (MacKenzie et al., 2014) suggested that isotope values for prey from the literature would be appropriate to use in the present study.

**Figure 4.**
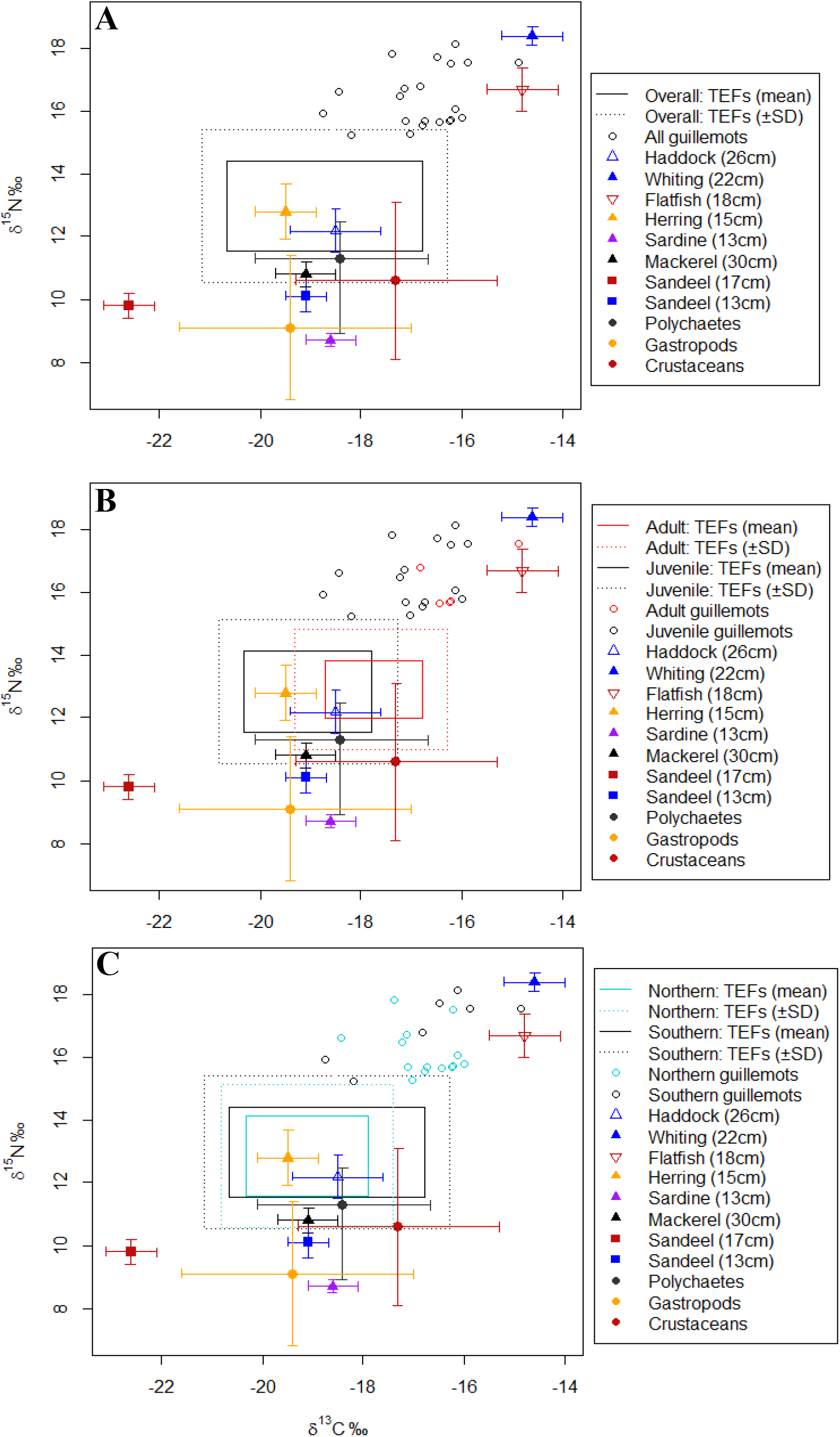
Carbon and nitrogen isotope values for guillemot secondary feathers, predicted prey and measured prey. (A) Overall dataset. (B) Guillemots split by age. (C) Guillemots split by location. Boxes show predicted ranges of prey isotope values of guillemots, using mean diet-feather trophic enrichment factors (TEFs) of 1.9‰ (δ^13^C) and 3.7‰ (δ^15^N) (solid boxes) and associated standard deviations of ±0.5‰ δ^13^C and ±1‰ δ^15^N (dotted boxes). Mean δ^13^C and δ^15^N values for several possible prey are also shown (from Käkelä et al. (2007) and Kürten et al. (2013); error bars show ±SD). Fish species are triangles and non-fish groups are filled circles (for scientific names see Appendix 6.6).

#### 2.6.iii. Euclidean methods to compare isotopic niche spaces

Euclidean methods to define and compare the isotopic niche space of species or groups within a community are widely used in stable isotope ecology (Layman et al., 2007). Such methods include Standard Ellipses and convex hulls. Conceptually, convex hulls are used to delineate the total extent of a species isotopic niche – in this case a two-dimensional niche space made up of δ^13^C and δ^15^N values. Standard Ellipses show the isotopic niche space occupied by a species or group to a given level of confidence (such as 95% confidence intervals). Whilst convex hulls crudely define isotopic niche space and measures its area subject to sampling biases and sensitivity to sample sizes, Standard Ellipses are more robust. These ellipses are estimated using Bayesian inference and incorporate uncertainty from sampling bias and different sample sizes into metrics of niche space and area (Jackson et al., 2011).

Therefore, differences in the two-dimensional δ^13^C and δ^15^N isotopic space occupied by juvenile and adult birds, as well as by northern and southern birds, were investigated using the SIBER package in R (Stable Isotope Bayesian Ellipses in R; Jackson et al., 2011). This approach utilises a statistical routine similar to bootstrapping, assigning measures of uncertainty iteratively using Markov-Chain Monte Carlo (MCMC) simulations to construct ellipse parameters. The most likely ellipse estimated using this Bayesian methodology can then be plotted (Bayesian Standard Ellipses corrected for small sample sizes by two degrees of freedom, (n-1) per axis; SEA_c_) over scatterplots of δ^13^C and δ^15^N values for birds. 95% confidence interval bivariate ellipses were plotted in all cases, giving a relatively robust representation of niche occupied by different groups of birds. Niche area (‰^2^) and overlap between the niche spaces was calculated through making posterior draws (100, 000) of SEA_c_ parameters. Using bivariate isotopic data for all individual birds, the isotopic variation in δ^13^C and δ^15^N values were incorporated into the posterior distribution, allowing valid comparisons in niche space and area between these different age classes and recovery locations (Jackson et al., 2011). Bivariate normality assumptions held under Mardia’s Multivariate Normality Test (Korkmaz et al., 2016). Investigating whether there were differences between age classes within southern and northern birds was not possible due to limited sample sizes of adults (SIBER analysis requires a minimum of five observations).

## 3. Results

Isotopic data were analysed from 21 individual birds including adults and juveniles from northern and southern recovery locations (Table 2).

**Table 2.** Mean δ^13^C and δ^15^N values (±SD) from feathers grown during the post-breeding moult of northern and southern guillemots.

| | $\delta^{13}\text{C}$ (‰) | | | | | | $\delta^{15}\text{N}$ (‰) | | | | | |
| --- | --- | --- | --- | --- | --- | --- | --- | --- | --- | --- | --- | --- |
|  | Juvenile | n | Adult | n | Overall | n | Juvenile | n | Adult | n | Overall | n |
| **Northern** | -16.92<br>$\pm 0.68$ | 11 | -16.29<br>$\pm 0.13$ | 3 | -16.78<br>$\pm 0.66$ | 14 | 16.31<br>$\pm 0.82$ | 11 | 15.70<br>$\pm 0.03$ | 3 | 16.18<br>$\pm 0.77$ | 14 |
| **Southern** | -17.08<br>$\pm 1.30$ | 5 | -15.85<br>$\pm 1.38$ | 2 | -16.73<br>$\pm 1.34$ | 7 | 16.92<br>$\pm 1.26$ | 5 | 17.18<br>$\pm 0.52$ | 2 | 16.99<br>$\pm 1.05$ | 7 |
| **Overall** | -16.97<br>$\pm 0.88$ | 16 | -16.12<br>$\pm 0.73$ | 5 | -16.77<br>$\pm 0.91$ | 21 | 16.50<br>$\pm 0.98$ | 16 | 16.29<br>$\pm 0.85$ | 5 | 16.45<br>$\pm 0.93$ | 21 |

### 3.1. Likely foraging areas of adults and juveniles

Firstly, care must be taken in interpreting these foraging area maps as they show the likeliest places where guillemots may have foraged in the past. Therefore, when considering, for example, adult birds overall (Fig.2A), spots of dark blue correspond to where the most likely distributions of all five birds within that sample overlap. Overall, adults and juveniles showed very similar foraging areas during the post-breeding moult (Fig.2A, B). Most likely foraging locations in both adults and juveniles appeared as a band along the coast of the eastern UK, stretching from slightly north of the Humber Estuary up to Orkney, as well as two patches in the southern and eastern North Sea: 1.) off the coast of western Denmark and southern Norway and 2.) slightly north of The Netherlands. However, juveniles did show wider likely foraging areas than adults, stretching between these patches (Fig.2B) – particularly evident in southern birds (Fig.2E, F). However, this is probably because of low sample sizes for adult birds, resulting in a slightly more restricted range of likely foraging areas. Some of these areas for both ages were also found in limited parts of the central North Sea.

Age classes did not show any significant isotopic differences overall or within subsets of northern or southern birds (p>0.05 in all cases; Table 3). However, large parameter estimates for the effect of age, particularly in δ^13^C values, appeared in the overall dataset: mean δ^13^C values in adult feathers were 0.81‰ (SE: ±0.48‰) greater compared to juveniles (t_19_=1.7, p=0.11; Table 3).

**Table 3.** General linear models (t-tests) for the whole dataset, as well as subsets for northern and southern birds. Significant differences (p<0.05) are denoted by asterisks (*). Parameter estimate (Est.) and standard error units are ‰.

| | Parameter | $\delta^{13}\text{C}$ | | | | | | $\delta^{15}\text{N}$ | | | | | |
| --- | --- | --- | --- | --- | --- | --- | --- | --- | --- | --- | --- | --- | --- |
|  |  | Est. | Std. error | df | Test stat. (t) | p-value | R<sup>2</sup> | Est. | Std. error | df | Test stat. (t) | p-value | R<sup>2</sup> |
| Northern | Age | 0.63 | 0.41 | 12 | 1.5 | 0.15 | 0.10 | -0.60 | 0.49 | 12 | 1.2 | 0.24 | 0.04 |
| Southern | Age | 1.23 | 1.10 | 5 | 1.1 | 0.31 | 0.04 | 0.26 | 0.96 | 5 | 0.3 | 0.80 | 0.18 |
| Overall | Age | 0.81 | 0.48 | 19 | 1.7 | 0.11 | 0.09 | -0.27 | 0.44 | 19 | 0.6 | 0.53 | 0.16 |
|  | Location | 0.03 | 0.43 | 19 | 0.1 | 0.95 | 0.08 | -0.97 | 0.40 | 19 | 2.4 | 0.03* | 0.20 |

Parameter estimates for increased mean δ^13^C values in adults compared to juveniles were also large in both northern and southern subsets: +0.63‰ (t_12_=1.5, p=0.15) and +1.23‰ (t_5_=1.1, p=0.31), respectively – although standard errors were also large (±0.41‰ and ±1.10‰, respectively; Table 3). When displayed graphically (red data points; Fig.3A), adults generally appeared further to the right-hand side of the x-axis (δ^13^C). This may suggest statistically significant differences were not detected due to small sample sizes. Nevertheless, these probably had little biological significance when considering the similarities in likely foraging areas (Fig.2). Conversely, large parameter estimates for reductions in mean δ^15^N values in adults compared to juveniles were reported within the northern subset (-0.60‰, t_12_=1.2, p=0.24; Table 3), albeit with large standard error (±0.49‰). These are less convincing when displayed graphically (adults are spread along the y-axis; Fig.3A). Low R^2^ values (Table 3) may also suggest inter-individual differences made up a large proportion of the variation in these data. This was also suggested by cluster analysis, where groupings do not split well by age (Fig.3B).

### 3.2. Likely foraging areas of northern and southern guillemots

There were also differences between northern and southern guillemots’ foraging areas (Fig.2C, D versus E, F). Northern guillemots were more likely to forage further north than southern guillemots, although there was overlap in foraging areas observed in the southern and eastern North Sea. However, the fact these southern birds were foraging in the North Sea is noteworthy considering they died foraging off the coast of southwestern England between December-February (Appendix 6.1, 6.3), suggesting they moved after the post-breeding moult between these ocean basins. Southern birds also had significantly higher δ^15^N values (+0.97‰ ± 0.40‰ than northern birds in the overall dataset (p<0.05, t_19_=2.4; Table 3) and cluster analysis resulted in one grouping containing only northern birds (bottom-right cluster; Fig.3B). There were no significant differences in δ^13^C between locations (p=0.95, t_19_=0.1; Table 3).

### 3.3. Diet

When considering guillemots overall, ranges of predicted isotope values for prey suggested they consumed haddock (a gadoid species) and herring (a clupeid species) (Fig.4A). With less certainty, considering standard deviations of measured and predicted prey isotope values, mackerel (another clupeid), polychaetes, gastropods, crustaceans and small sandeels (13cm) may also have contributed to a mixed diet (Fig.4A).

The range of prey isotope values for adults only included haddock, and possibly crustaceans and gastropods (considering their standard deviation), whilst juveniles may have consumed the full range of prey items described previously (Fig.4B). Considering standard deviation in estimated prey isotope values shown by dotted boxes (Fig.4B), adults may also have foraged on herring and mackerel. Therefore, there is not enough evidence to suggest juveniles and adults consume different prey during the post-breeding moult, although adults may consume a slightly less varied diet – although this could also be an artefact of small sample sizes. Adults and juveniles (pooled from both locations to ensure robust analysis) occupied slightly different isotopic niche space, mainly due to differences in δ^13^C values (as noted earlier, adults in red are shifted to the right; Fig.5A). There was slight overlap in 95% confidence interval ellipses in line with overlap in their likely foraging areas (Fig.2; Fig.5A). However, these niches were similarly sized which does not support the notion that juveniles had a wider range of likely foraging areas or diet (59% of MCMC simulated posterior ellipse areas for adults were larger than those for juveniles; Fig.5B). This is probably because SIBER analysis is robust against differences in sample sizes and incorporates these into ellipse areas.

**Figure 5.**
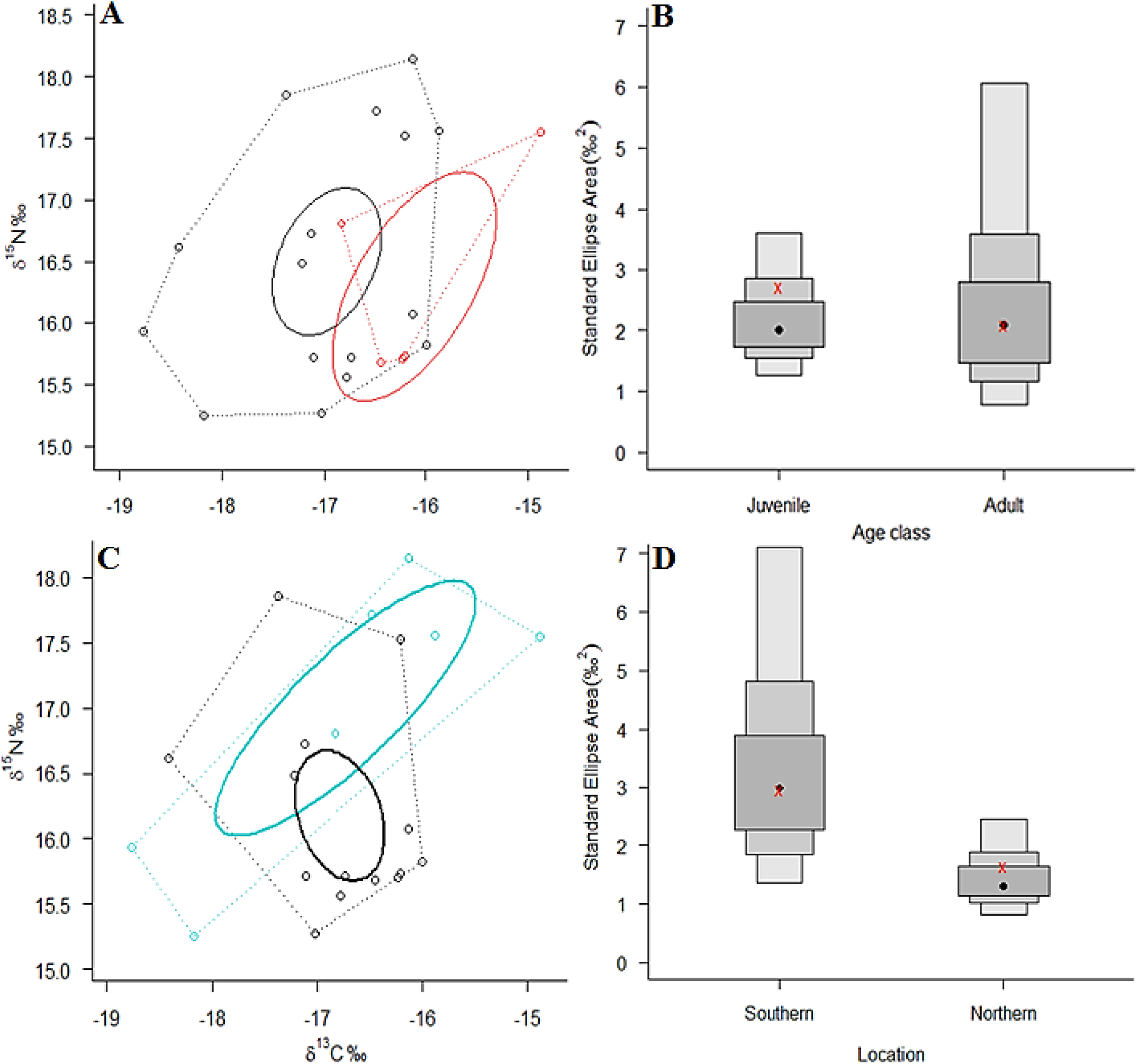
(A and C) Biplots of δ^13^C and δ^15^N values from feathers of guillemots. Black ellipse and black dotted lines show most likely 95% confidence ellipses and convex hulls, respectively, for juveniles (A) and northern birds (C). The equivalents in red are the same but for adult birds(A), whilst those in light blue are for southern birds (C). (B and D) Bayesian standard ellipse areas (SEA_c_, corrected for small samples in a bivariate distribution for 2df) for juveniles and adults (B) and southern and northern birds (D) based on 100 000 posterior draws of parameters. 50, 75 and 95% credibility intervals are shown from dark to light grey, respectively. Maximum likelihood estimates for SEAc are denoted by red x marks, whilst black dots denote modal maximum likelihood values for ellipse areas.

When considering isotopic niches of northern versus southern birds, there were differences due to δ^15^N values (Fig.5C), as demonstrated by the statistically significant difference detected by models (Table 3). However, there was overlap in 95% confidence ellipses, supporting the overlap observed in their foraging areas in the southern and eastern North Sea. Nevertheless, despite overlap in 95% credibility intervals (Fig.5D), northern birds occupied significantly smaller isotopic niche space than southern birds, possibly suggesting diet of southern birds was more limited (p<0.05, 2.9% of MCMC simulated posterior ellipse areas for southern birds were larger than those of northern birds; Fig.5C). A statistically smaller isotopic niche was probably detected because the range of predicted prey isotope values for northern birds fell inside the ranges of southern birds (Fig.4C). However, diet analysis suggested this finding carried negligible biological significance, since both northern and southern birds both foraged on the same broad range of fish and non-fish prey mentioned earlier (Fig.4C).

## 4. Discussion

The aims of this study were to investigate where guillemots forage and what they consume during the post-breeding moult, according to different age classes and recovery locations. Overall, isotopic analysis of guillemot feathers revealed that guillemots ranged widely throughout the western, eastern, and southern North Sea and had a mixed diet of fish and non-fish prey. Likely foraging areas of adults and juveniles were similar and there seemed to be few age-related differences in diet. Northern birds appeared to have a wider range of likely foraging areas, particularly in the northwestern North Sea, whilst both groups overlapped in the southern and eastern North Sea. Northern and southern birds both had similar diets despite these differences in the range of likely foraging areas. These findings highlight several areas for further discussion, such as why adult and juvenile foraging areas and diets were similar.

### 4.1. Similarities in adult and juvenile foraging areas and diet

There was conflicting evidence from the literature suggesting whether adult and juvenile foraging areas would be expected to be similar or different during the post-breeding moult. Observed similarities in their foraging areas may not only imply that adult guardians that accompany juveniles from their breeding colony remain with them during this time, but that non-guardian adults also travel to similar locations during the moult (Jones and Rees, 1985). Assertions by Harris and Swann (2002) that juveniles range more widely in the non-breeding season are poorly supported, however, this is relative to the breeding colony that birds are from which could not be determined in this study. Nevertheless, juveniles appeared to have wider ranges in their likely foraging areas, implying that they might also show more variation in where they forage after leaving a given colony. Another explanation bridging these differing ideas may be that juveniles and adults only remain together until adults return to their breeding colony around October (Wanless and Harris, 1986; Halley et al., 1995), after which juveniles disperse further over winter – as observed by Harris and Swann (2002). How the return of adults to their breeding colonies fits within the wider context of non-breeding season movements warrants further investigation.

Nevertheless, a point to acknowledge about the isoscape used by this study is that it is limited to the North Sea, central and eastern English Channel and minor parts of the western shores of the UK (Fig.1). Therefore, guillemots could have foraged in places with similar isotopic compositions outside the study area. The Bay of Biscay is one potential area as Le Rest et al. (2016) claimed a major proportion of British guillemots overwinter here; particularly juveniles and immature birds. However, Le Rest et al. (2016) considered winter distributions and it is possible that adults and juveniles forage in different places during the post-breeding moult to where they overwinter. Their assumption that most guillemots were ‘British’ was also based on the sheer number of overwintering birds, which seems invalid given there was no other information on breeding colonies. It is more likely these aggregations were part of the wider Northeast Atlantic and North Sea meta-population. The expansion of this isoscape (currently underway; C Trueman, 2017, personal communication, 30^th^ March), as well as increased sample sizes would enable a more conclusive investigation into whether different age classes of guillemots forage in different places during the post-breeding moult compared to winter. Furthermore, given adult guardians are usually males, an investigation into sex-related differences in foraging areas during the post-breeding moult and non-breeding season would also be useful; such differences have been found in Northern Fulmars (*Fulmarus glacialis*) during the pre-laying exodus (Edwards et al., 2016).

There seemed to be few age-related differences in diet, as both adults and juveniles probably fed on a mixture of gadoids, clupeids and possibly polychaetes, gastropods, crustaceans and small sandeels. This is most likely because adult guardians stay with juveniles during the post-breeding moult and feed them prey (Jones and Rees, 1985). Although the dietary proportions of each prey item could not be inferred with certainty from these analyses, the great similarities in diet do not seem to support the age-related dietary preferences observed by Lorentsen and Anker-Nilssen (1999) in winter. This may suggest that after adults return to colonies in October (Wanless and Harris, 1986; Halley et al., 1995), juvenile diet may be restricted to more catchable prey by their lack of experience and swimming ability (Lorentsen and Anker-Nilssen, 1999).

### 4.2. Diet and foraging areas of guillemots recovered from different locations

Southern birds had significantly higher δ^15^N values than northern birds, suggesting either that they foraged in similar places but on different prey, or that these guillemots foraged in different places with different isotopic compositions. A lack of differences in diet between northern and southern guillemots appeared to suggest that the geographical differences in likely foraging areas caused these differences in δ^15^N values. These findings imply southern guillemots may adopt different movement strategies to northern guillemots during the non-breeding season. Between the post-breeding moult and when birds were recovered in winter, southern birds seemed to move southwest-wards to Cornish and Celtic Sea fisheries, whilst northern birds either moved northwest-ward or remained in the northwestern North Sea. Although this assumes birds moved in a straight line, these inferred movements provide a basis for further investigation of movement strategies during the non-breeding season.

As mentioned previously, the restricted area considered by this isoscape means that southern guillemots could have foraged in the Celtic Sea during the post-breeding moult. However, there was no evidence they foraged in the central and eastern English Channel during this period, so it is more likely that these birds either solely forage in the Celtic Sea and western English Channel, or further east in the North Sea. A more comprehensive study could build on these findings and uncertainties by considering birds from known colonies around the UK. This would enable a thorough assessment of the overlap in foraging areas between different UK breeding colonies to better understand guillemot movements during the non-breeding season. Using GLS tagging, McFarlane Tranquilla et al. (2014) found two major winter movement strategies amongst individuals from different colonies in the Northwest Atlantic: inshore versus offshore. Over three to four years, these guillemots either kept the same strategy or were flexible between strategies, suggesting long-term research may be needed to fully understand movement strategies in the North Sea (McFarlane Tranquilla et al., 2014).

Another point to consider when comparing northern and southern birds is their different levels of sampling bias. Southern birds were thought to be less biased samples, since birds are likely to have been healthy before being bycaught and therefore were more likely to represent at-sea distributions of guillemots (Peterz and Blomqvist, 2010). Nevertheless, linear models suggested both northern and southern subsets showed similar patterns in age-related differences. Moreover, the likely foraging areas of northern and southern birds during the post-breeding moult are consistent with those reported in northern Scotland (Wright and Begg, 1997) and more widely through GLS tagging (Harris et al., 2015). This suggests guillemots from both recovery locations were more representative than expected, despite small sample sizes and possible biases.

Nevertheless, without a more comprehensive study, the most valid way to draw rigorous conclusions about wider guillemot populations in the UK is to consider the overall foraging areas reported by this study. These closely matched the distribution of guillemots reported in Harris et al. (2015) within the North Sea during the post-breeding moult. Additionally, the substantial intracolonial variation in moulting locations observed by Harris et al. (2015) also suggests inter-individual differences constitute a major portion of the variation in moulting locations. This may explain why most model fits were poor when testing for isotopic differences between ages and recovery locations. This would mean that different colonies in the UK may show great overlap in foraging areas during the moult, although this requires further investigation as these results were from a single colony.

Results from Harris et al. (2015) also indicate that an expansion of the isoscape is necessary, since guillemots moved into areas outside of the North Sea (the Barents Sea and Kattegat) during the post-breeding moult. Nevertheless, the consistency between the findings of this study and other studies from the North Sea (Wright and Begg, 1997; Harris et al., 2016) shows that although accuracy for feather assignments was predicted to be approximately 75%, this isoscape provided relatively robust assignments. This is also noteworthy considering the small sample sizes available and the many assumptions this method relies on when geographically assigning mobile animals from higher trophic levels (Trueman et al., 2016).

### 4.3. Considering diet and foraging areas in the context of potential threats

Wright and Begg (1997) suggested depth and tidal current rate were also important factors in determining moulting distributions, as they influence guillemot feeding efficiency and habitat suitability for their prey. Further investigation of correlations between guillemot and prey distribution during the post-breeding moult is needed to understand how important prey-based drivers of guillemot distribution are. However, these will be difficult to disentangle considering the wide range of fish and non-fish prey that guillemots consumed – as expected, since the post-breeding moult is a time of major dietary change between breeding season and non-breeding season diets (Blake et al., 1985; Lorentsen and Anker-Nilssen, 1999; Sonntag and Hüppop, 2005). This mixed guillemot diet (including clupeids and gadoids, crustaceans, polychaete worms, gastropods and possibly small sandeels) also supports the idea that these dietary changes also occur in more southerly parts of the North Sea – previously these changes had only been reported in the northern North Sea (Blake et al., 1985). No appropriate data were available for some possible prey, such as cod and pollack or pipefish and gobies reported in winter diet by Sonntag and Hüppop (2005), so these findings do not preclude these from the diet of guillemots during the post-breeding moult.

Interestingly, most of the fisheries where southern guillemots were recovered during the winter targeted gadoid species (cod, pollack and haddock; Appendix 6.3), supporting assertions by Blake et al. (1985) that guillemots continue to prey on these species in winter. These observations also highlight how the diet of guillemots may bring them into conflict with fisheries and the importance of studies considering diet outside of the breeding season in assessing the risks fisheries pose to seabird populations. The likely foraging areas identified by this study substantially overlap with gillnet fishing effort, particularly in the southern and eastern North Sea (STECF, 2016). Therefore, based on these findings, attempts to mitigate bycatch should focus on these areas of greatest risk for guillemots, particularly gadoid and clupeid gillnet fisheries. As large aggregations of guillemots are likely to use these foraging areas, major pollution events such as oil spills are also major threats to their populations during this time (Le Rest et al., 2016). Concerningly, high levels of shipping traffic also overlap with foraging areas in the southern and eastern North Sea (Marine Traffic, 2017), putting guillemots at great risk from pollution caused by shipping accidents. Moreover, offshore oil installations also overlap with likely foraging areas of guillemots in the southern North Sea (particularly above The Netherlands; EU Offshore Authorities Group, 2012) also putting guillemots at further risk of more substantial pipeline oil spills. Overall, likely foraging areas of guillemots during the post-breeding moult seem to be highly threatened by anthropogenic activities.

### 4.4. Conclusions and future research

In summary, this study demonstrated that Common Guillemots do not show convincing age-related differences in their diet or foraging areas during the post-breeding moult. Further evidence has also been found that seasonal dietary changes reported by Blake et al. (1985) also occur in guillemots in the southern North Sea. Whilst diet did not vary between guillemots recovered from different locations in the UK, their use of alternative foraging areas to one another implied they exhibited different movement strategies during the non-breeding season. The consistency between this study’s findings and the limited literature on guillemot movements in the North Sea during the post-breeding moult also suggested that this isoscape was a relatively robust method to geographically assign highly mobile animals from higher trophic levels. Therefore, this isoscape has the potential to help uncover more about the spatial ecology of guillemots and other seabirds during the non-breeding season.

To build on these findings, studies combining an expanded isoscape study area with GLS tagging should focus on whether sex and colony are important factors in determining guillemot movements during the non-breeding season. Answering these questions will give a more complete understanding of guillemot movement strategies during the non-breeding season and help discern effective ways to protect UK breeding colonies from anthropogenic threats. This is of paramount importance given that guillemot foraging areas were shown to be highly vulnerable to gillnet fishing and oil spills, particularly in the southern and eastern North Sea. Efforts to conserve guillemots in the North Sea should focus on these regions to reduce the risk and possible impacts of these threats.

## 6. Acknowledgements

My thanks go to Dr Simon Northridge, Dr Nora Hanson, Professor Michael Harris, Martin Heubeck, Angus Calder, the University of St Andrews Stable Isotope Laboratory, Jason Newton, SUERC, Dr Clive Trueman, Katie St John Glew, Allen Kingston, all fisheries bservers involved in recovering guillemot carcasses from fisheries off southwestern England, and Grania Smith.

## 7. Appendix

**6.1.**
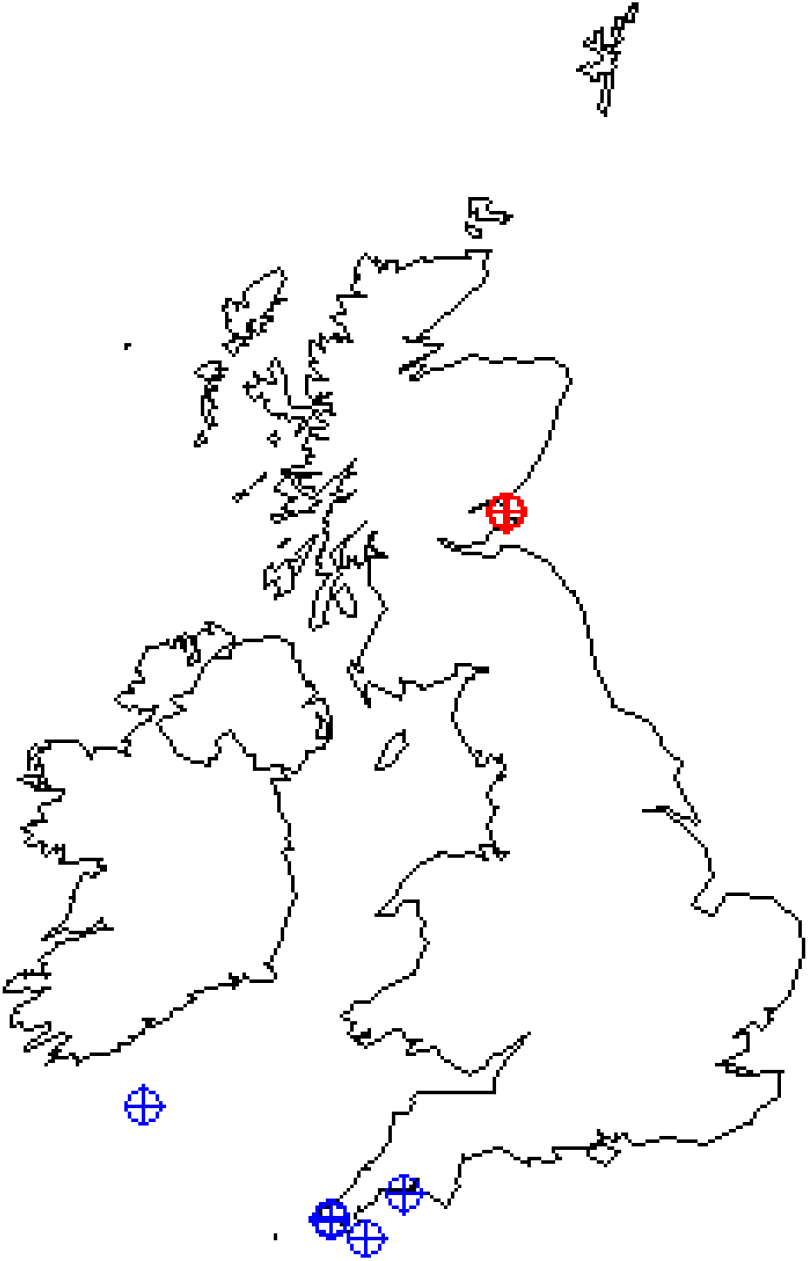
Map of the British Isles showing where Common Guillemot carcasses were recovered from – red points refer to birds recovered from beaches in St Andrews, Scotland, whilst blue points show birds recovered from mixed static net fisheries in Cornish waters and the adjacent Celtic Sea (see Appendix 6.3 for details).

**6.2.** Data for all individual birds, including their recovery dates, their feathers’ mean δ^15^N and δ^13^C values, their age (J=juvenile, A=adult) and their recovery location (N=northern (St Andrews, Scotland), S=southern (Mixed static net Cornish and Celtic Sea fisheries)).

| ID | Date | Mean $\delta^{13}\text{C}$ | Mean $\delta^{15}\text{N}$ | Age | Location |
| --- | --- | --- | --- | --- | --- |
| 6 | 09/10/2016 | -16.44 | 15.68 | A | N |
| 7 | 09/10/2016 | -18.42 | 16.62 | J | N |
| 8 | 09/10/2016 | -17.11 | 15.71 | J | N |
| 9 | 09/10/2016 | -16.20 | 15.73 | A | N |
| 10 | 09/10/2016 | -16.13 | 16.07 | J | N |
| 17 | 12/10/2016 | -16.73 | 15.72 | J | N |
| 18 | 11/10/2016 | -17.22 | 16.49 | J | N |
| 19 | 11/10/2016 | -17.12 | 16.73 | J | N |
| 21 | 12/10/2016 | -16.21 | 17.52 | J | N |
| 23 | 12/10/2016 | -16.23 | 15.70 | A | N |
| 24 | 12/10/2016 | -16.78 | 15.56 | J | N |
| 25 | 12/10/2016 | -17.38 | 17.85 | J | N |
| 33 | 25/10/2016 | -16.00 | 15.82 | J | N |
| 35 | 18/02/2017 | -17.03 | 15.27 | J | N |
| 41 | 22-23/01/2017 | -16.13 | 18.14 | J | S |
| 42 | 22/01/2017 | -16.83 | 16.81 | A | S |
| 43 | 17/12/2016 | -18.76 | 15.93 | J | S |
| 44 | 01-06/12/2016 | -16.48 | 17.72 | J | S |
| 45 | 06/01/2017 | -14.88 | 17.55 | A | S |
| 46 | 17/02/2017 | -18.17 | 15.25 | J | S |
| 47 | 17/02/2017 | -15.87 | 17.56 | J | S |

**6.3.** Details of fisheries guillemots were recovered from, along with other information on the type of net used, target of the fishery and coordinates of the static nets.

| ID | Trip | Haul no. | Lat. | Long. | Net type | Target |
| --- | --- | --- | --- | --- | --- | --- |
| 41 | JH282 | 3 | 50.08083 | -5.47750 | gill | pollack, cod |
| 42 | JH282 | 4 | 50.08267 | -5.46700 | gill | pollack, cod |
| 43 | JH280 | 4 | 50.10050 | -5.48917 | gill | haddock |
| 44 | JH278 | 5 | 51.09200 | -8.32750 | gill | hake |
| 45 | AK514 | 1 | 49.92033 | -4.92983 | tangle | anglerfish |
| 46 | PJS704 | 1 | 50.31117 | -4.34917 | gill | pollack |
| 47 | PJS704 | 1 | 50.31133 | -4.34933 | gill | pollack |

**6.4.**
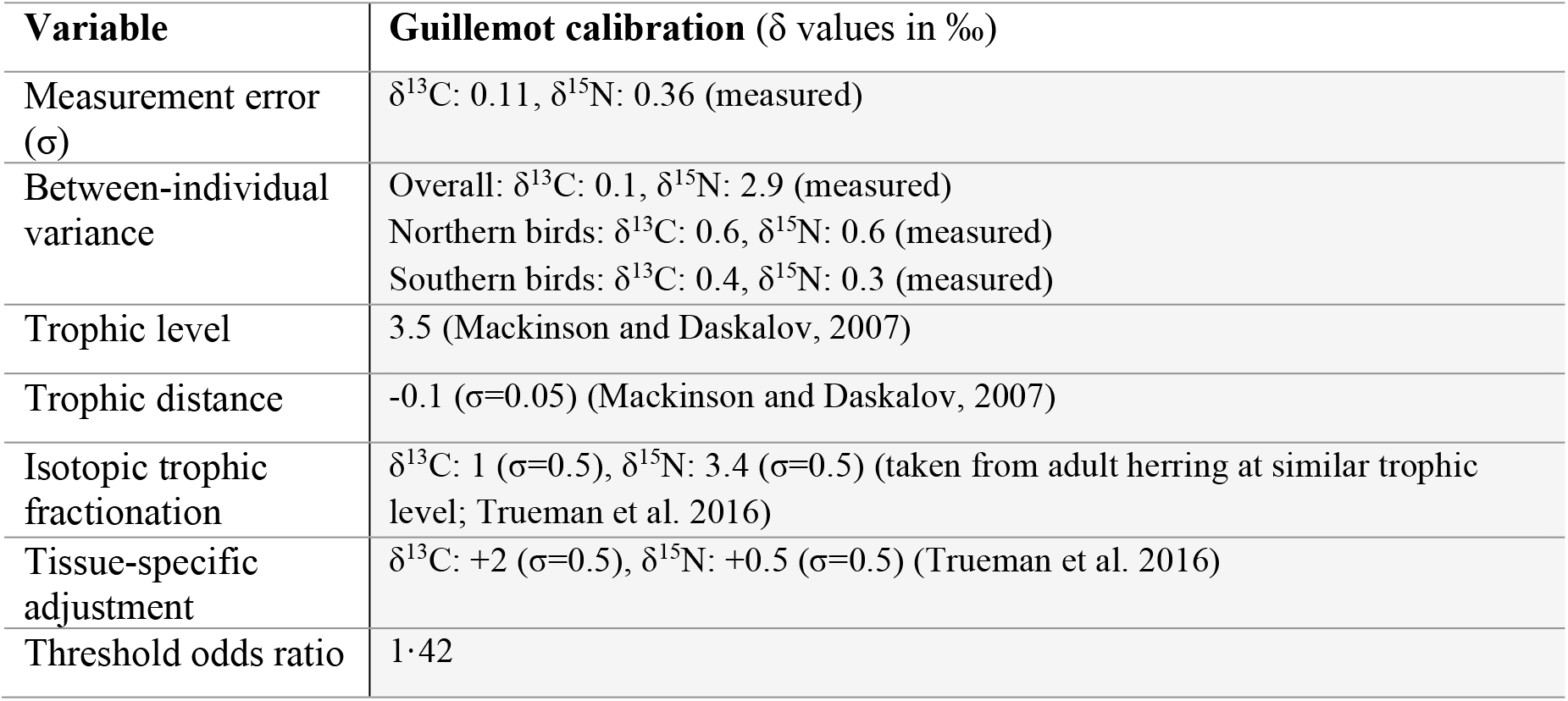
Assignment conditions for the stable isotope-based geographic assignment of common guillemots to isoscapes derived from jellyfish tissues

| Variable | Guillemot calibration ($\delta$ values in ‰) |
| --- | --- |
| Measurement error ($\sigma$) | $\delta^{13}\text{C}$: 0.11, $\delta^{15}\text{N}$: 0.36 (measured) |
| Between-individual variance | Overall: $\delta^{13}\text{C}$: 0.1, $\delta^{15}\text{N}$: 2.9 (measured)<br>Northern birds: $\delta^{13}\text{C}$: 0.6, $\delta^{15}\text{N}$: 0.6 (measured)<br>Southern birds: $\delta^{13}\text{C}$: 0.4, $\delta^{15}\text{N}$: 0.3 (measured) |
| Trophic level | 3.5 (Mackinson and Daskalov, 2007) |
| Trophic distance | -0.1 ($\sigma=0.05$) (Mackinson and Daskalov, 2007) |
| Isotopic trophic fractionation | $\delta^{13}\text{C}$: 1 ($\sigma=0.5$), $\delta^{15}\text{N}$: 3.4 ($\sigma=0.5$) (taken from adult herring at similar trophic level; Trueman et al. 2016) |
| Tissue-specific adjustment | $\delta^{13}\text{C}$: +2 ($\sigma=0.5$), $\delta^{15}\text{N}$: +0.5 ($\sigma=0.5$) (Trueman et al. 2016) |
| Threshold odds ratio | 1.42 |

**6.5.**
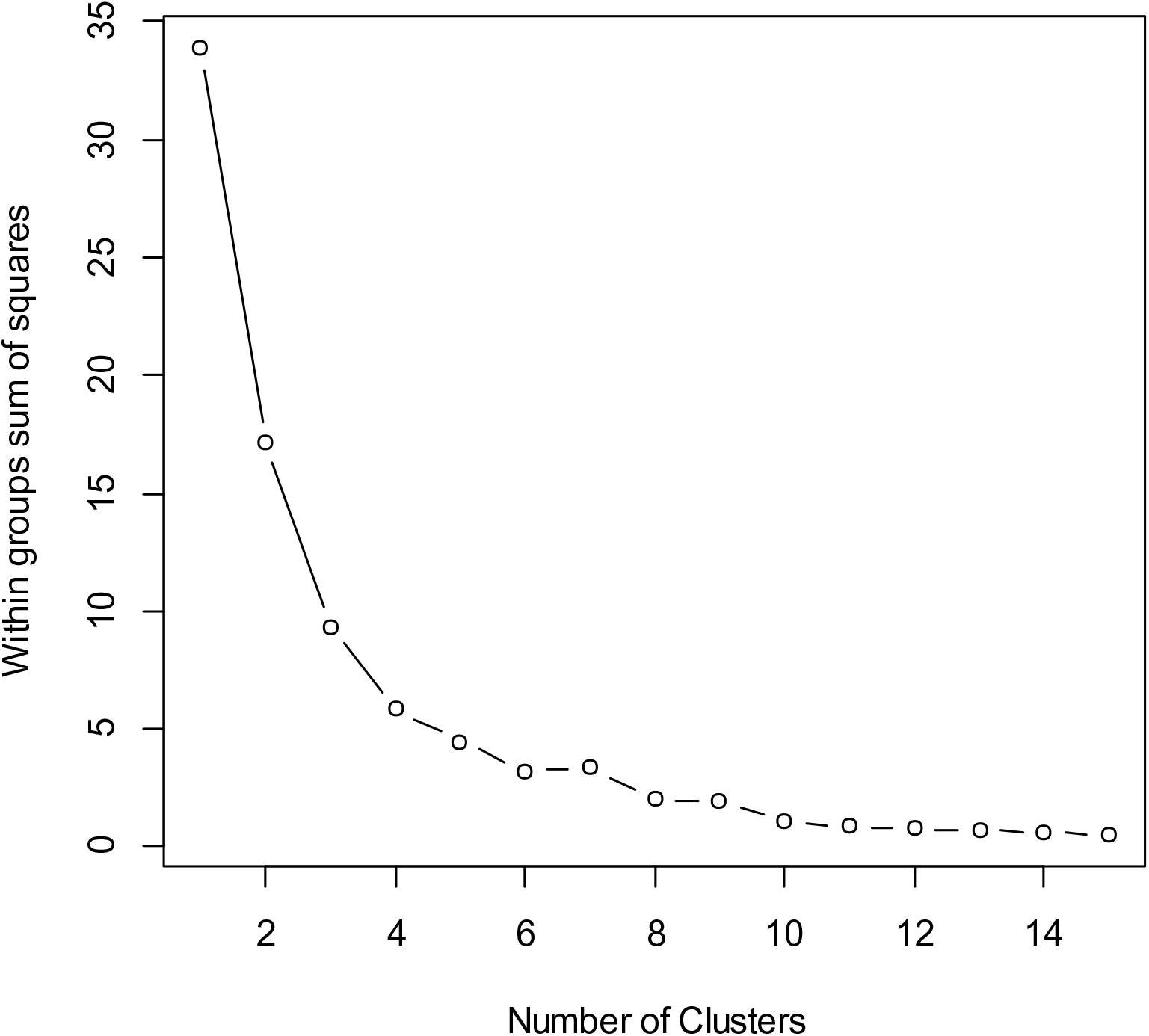
A plot of the total within-groups sums of squares against the number of clusters in a K-means solution for the whole bivariate dataset of δ^15^N and δ^13^C values. This plot can identify the best number of clusters that this cluster analysis method should use to find the most cohesive clusters from the available data. A levelling off after four clusters suggests this is the most appropriate number for cluster analysis to assign to the data.

**6.6.** Summary of mean δ^15^N and δ^13^C values (±SD) for all possible prey, with common and scientific names, considered in this study. ^1^ = data from Käkelä et al. (2007). ^2^ = data from Kürten et al. (2013).

| Common name | Scientific name | $\delta^{13}\text{C}$ (‰) | $\delta^{15}\text{N}$ (‰) |
| --- | --- | --- | --- |
| Haddock<sup>1</sup> | *Melanogrammus aeglefinus* | $-18.5 \pm 0.9$ | $12.2 \pm 0.7$ |
| Whiting<sup>1</sup> | *Merlangius merlangus* | $-14.6 \pm 0.6$ | $18.4 \pm 0.3$ |
| Flatfish<sup>1</sup> | Flatfish spp. | $-14.8 \pm 0.7$ | $16.7 \pm 0.7$ |
| Herring<sup>1</sup> | *Clupea harengus* | $-19.5 \pm 0.6$ | $12.8 \pm 0.9$ |
| Sardine<sup>1</sup> | *Sardina pilchardus* | $-18.6 \pm 0.5$ | $8.7 \pm 0.2$ |
| Mackerel<sup>1</sup> | *Scomber scombrus* | $-19.1 \pm 0.6$ | $10.8 \pm 0.4$ |
| Large sandeel (17cm) <sup>1</sup> | *Ammodytes marinus* | $-22.6 \pm 0.5$ | $9.8 \pm 0.4$ |
| Small sandeel (13cm) <sup>1</sup> | *Ammodytes marinus* | $-19.1 \pm 0.4$ | $10.1 \pm 0.5$ |
| Crustaceans<sup>2</sup> | Crustacea | $-19.3 \pm 2.3$ | $9.2 \pm 2.3$ |
| Gastropods<sup>2</sup> | Gastropoda | $-17.3 \pm 2.0$ | $10.6 \pm 2.5$ |
| Polychaetes<sup>2</sup> | Polychaeta | $-18.4 \pm 1.7$ | $11.2 \pm 2.3$ |

